# Bacterial supergroup specific “Cost” of *Wolbachia* infections in *Nasonia vitripennis*

**DOI:** 10.1101/2021.09.10.459769

**Authors:** Alok Tiwary, Rahul Babu, Ruchira Sen, Rhitoban Raychoudhury

**Affiliations:** Department of Biological Sciences, Indian Institute of Science Education and Research (IISER) Mohali, Knowledge City, Sector 81, SAS Nagar, Manauli, PO 140306, Punjab, India; Zoological Survey of India, Kolkata, West Bengal-700053, India; Guru Gobind Singh College, Sector 26, Chandigarh-160019, India

**Keywords:** *Wolbachia*, *Nasonia*, life-history traits, Quantitative PCR, Mann-Whitney U, Prospermatogeny, Cytoplasmic incompatibility

## Abstract

The maternally-inherited endosymbiont, *Wolbachia*, is known to alter the reproductive biology of its arthropod hosts for its benefit and can induce both positive and negative fitness effects in many hosts. Here we describe the effects of the maintenance of two distinct *Wolbachia* infections, one each from supergroups A and B, on the parasitoid host *Nasonia vitripennis*. We compare the effect of *Wolbachia* infections on various traits between the uninfected, single A infected, single B infected, and the double infected strains with their cured versions. Contrary to the previous reports, our results suggest that there is a significant cost associated with the maintenance of *Wolbachia* infections where traits like family size, fecundity, longevity, and rates of male copulation are compromised in *Wolbachia* infected strains. The double infected and supergroup B infection strains show higher *Wolbachia* titer than supergroup A. The double infected *Wolbachia* strain has the most detrimental impact on the host as compared to single infections. Moreover, there is a supergroup-specific negative impact on these wasps as the supergroup B infections elicit the most pronounced negative effects. These findings raise important questions on the mechanism of survival and maintenance of these reproductive parasites in arthropod hosts.

## Introduction

*Wolbachia* are maternally-inherited, obligatory intracellular endosymbionts of the order Rickettsiales (Hertig et al., 1927) that are widely found in arthropods and filarial nematodes (Bandi et al. 1992; Rousset et al. 1992; Weinert et al., 2015). These bacteria often alter host reproductive biology with mechanisms like male-killing, feminization, parthenogenesis, and cytoplasmic incompatibility (CI), to enhance their own transmission (Werren et al., 2008). While CI leads to an increase in the number of infected individuals in the population, male-killing, and feminization shifts the offspring sex ratio towards females, which is the transmitting sex for *Wolbachia*. Thus, *Wolbachia* increases the fitness of the infected hosts as it increases its own rate of transmission. While these cases highlight a parasitic effect of *Wolbachia*, there are several examples where no such effect is discernible (Hoffmann et al., 1996). Moreover, there are also examples where *Wolbachia* has now become a mutualist and offers specific and quantifiable benefits to its host. One such example of an obligate mutualism with *Wolbachia* has been reported in *Asobara tabida* where females cured of *Wolbachia* are unable to produce mature oocytes (Dedeine et al., 2001). Such examples of arthropod-*Wolbachia* mutualism have now been reported from various arthropod taxa (Pike & Kingcombe, 2009; Miller et al., 2010). This shift from parasitic to mutualistic effect can also happen in facultative associations as seen in *Drosophila simulans*, where within a span of just two decades, *Wolbachia* has evolved from a parasite to a mutualist (Weeks et al., 2007). However, for the majority of *Wolbachia*-host associations, no such reports of mutualistic benefits exist and instead reveal many negative effects on the host. In addition to reproductive traits, many other life-history traits like longevity and developmental time are also known to be compromised. A review of such negative effects of *Wolbachia* on hosts where CI is prevalent is presented in Table 1. In *Trichogramma kaykai and T. deion*, the infected (thelytokous) line shows reduced fecundity and adult emergence rates than the antibiotically cured (arrhenotokous) lines (Hohmann et al., 2001; Tagami et al., 2001). *Leptopilina heterotoma*, a *Drosophila* parasitoid, has adult survival rates, fecundity, and locomotor performance, of both sexes, severely compromised in *Wolbachia* infected strains (Fleury et al., 2000). Preimaginal disc mortality has been observed in both sexes of insecticide-resistant *Wolbachia*-infected strains of *Culex pipiens* (Duron et al., 2006). *Wolbachia* infections can also result in a range of behavioral changes and altered phenotypes in *Aedes aegypti* (Turley et al., 2009).

**Table 1.** Negative fitness effects of CI-inducing *Wolbachia*.

| *Wolbachia* Supergroup | Genera | Species | *Wolbachia* Strain | Host Sex | Negative effect | Reference |
| --- | --- | --- | --- | --- | --- | --- |
| A | *Drosophila* | *D. melanogaster* | A-wMelPop | Female/Male | Tissue degeneration, reduced lifespan | Min et al. 1997; Reynolds et al. 2003 |
|  |  |  | A-wMelPop | Female/Male | Decreased response to food cues | Peng et al. 2008 |
|  |  |  | A-wMel | Female | Reduced body size | Hoffmann et al. 1997 |
|  |  |  | A-wMel | Female | Reduced fecundity after a dormancy period | Kriesner et al. 2016 |
|  |  | *D. simulans* | A-wRi | Female | Reduction in fecundity | Hoffmann et al., 1988, 1990 |
|  |  |  | A-wRi | Female | Reduction in fecundity | Snook et al. 2000 |
|  |  |  | A-wRi | Male | Lesser sperm cysts, reduced fertility | Snook et al. 2000 |
|  |  |  | A-wHa | Female | Reduction in fecundity | Fytrou et al. 2006 |
|  |  |  | A-wHa | Female/Male | Reduced thorax length, reduction in an immune response against parasitoid infection | Fytrou et al. 2006 |
|  |  |  | A-wRi | Male | Reduced sperm competition | Crespigny et al. 2006 |
|  |  | *D. sukuzii* | A-wSuz | Female | Reduced progeny family size | Hamm et al. 2014 |
| B | *Aedes* | *A. albopictus* | A-wAlb, B-wAlb | Female | Reduced lifespan, reduction in fecundity | Islam et al. 2006; Sun et al. 2009 |
|  |  | *A. aegypti* | A-wMelPop | Female | Reduced lifespan, reduction in fecundity | Ross et al. 2019 |
|  |  |  | A-wMelPop | Female/Male | Reduced lifespan | Mcmeniman et al. 2009 |
|  |  |  | A-wMelPop | Female | Reduction in fecundity, reduced blood-feeding success | Turley et al. 2009, 2013; Allman et al. 2020 |
|  | *Culex* | *C. pipiens* | B-wPip | Female/Male | Embryonic Mortality | Duron et al. 2005 |
|  |  | *C. quinquefasciatus* | B-wPip | Female | Reduced fecundity | Almeida et al. 2011 |

These negative effects of *Wolbachia* on their hosts are not unexpected. The presence of bacteria within a host entails sharing of nutritional and other physiological resources (Kobayashi & Crouch, 2009; Whittle et al., 2021), especially with *Wolbachia* as they are obligate endosymbionts and cannot survive without cellular resources derived from their hosts (Foster et al., 2005., Slatko et al., 2010). For example, *Wolbachia* is known to compete with the host for key resources like cholesterol and amino acids in *A. aegypti* (Caragata et al., 2014). The precise molecular mechanisms of many of these negative effects have not been ascertained and are generally ascribed to partitioning off of host nutrients for its benefit, but what is clear is that *Wolbachia* infections can impose severe nutritional demands on their hosts (Ponton et al., 2014). However, it is also known that *Wolbachia* can elicit antipathogenic responses from their hosts where the host resistance or tolerance to the infection increases (Zug & Hammerstein, 2015). For example, *Wolbachia* can utilize the Immune deficiency (IMD) and Toll pathways (Pan et al., 2018) and increased ROS (Reactive Oxygen Species) levels in *Wolbachia*-transfected *A. aegypti* mosquitoes, inhibiting the proliferation of the Dengue virus (Pan et al., 2012) as well as against *Drosophila* C viruses (Teixeira et al., 2008). Such immune responses require additional allocation of resources, which can further affect other physiological traits of the host. This concept of a “cost of immunity” is well-established and suggests a trade-off between immunity and other life-history traits (Zuk & Stoehr, 2002). For example, elevated ROS levels negatively affect many host traits like longevity, and fecundity (Dowling & Simmons, 2009; Monaghan et al., 2009; Selman et al., 2012; Moné et al., 2014). Thus, there is sufficient evidence to conclude that *Wolbachia* can have substantial negative effects on the overall fitness of its host.

In this study, we investigate what, if any, are the negative effects on the wasp *N. vitripennis* due to its two resident CI inducing *Wolbachia* infections? *N. vitripennis*, being cosmopolitan, has been used to study *Wolbachia* distribution, acquisition, spread, and *Wolbachia*-induced reproductive manipulations (Werren et al., 2008; Landmann, 2019). However, the effect of the endosymbiont on the host life-history traits of this wasp remains poorly understood with conflicting reports. In some *N. vitripennis* strains, *Wolbachia* was reported to be the cause of enhanced fecundity (Stolk & Stouthamer, 1996), but a similar effect was not observed in some other strains (Bordenstein & Werren, 2000). *N. vitripennis* harbor two *Wolbachia* supergroup infections, one each from supergroup A and supergroup B (Perrot-Minnot et al., 1996), and the presence of these two infections have been found in all strains of *N. vitripennis* from continental North America to Europe (Raychoudhury et al., 2010), indicating that it has reached fixation across the distribution of its host. The two *Wolbachia* in *N. vitripennis* together cause complete CI, but single infections of supergroup A *Wolbachia* causes incomplete CI while supergroup B infections still show complete CI (Perrot-Minnot et al., 1996). In this study, we investigate the effects of *Wolbachia* infections in a new strain of *N. vitripennis* recently obtained from the field. This strain, like other *N. vitripennis* strains, has two *Wolbachia* infections, one each from the supergroup A and B. Sequencing of the five alleles from the well-established multi-locus strain typing system (Baldo et al., 2006) reveal no sequence dissimilarity across the two infections (Prazapati, personal communication) indicating this new strain is also infected by the same or very similar *Wolbachia* strains that are present across the distribution of *N. vitripennis* (Raychoudhury et al., 2010b). To compare supergroup specific effects, these two infections have been separated into single infected strains. A comparative analysis between the double infected, supergroup A infected, supergroup B infected and uninfected strains revealed a consistent pattern of decreased longevity, quicker sperm depletion and reduced family size for infected individuals. While supergroup B infection has a more pronounced negative effect on most of the traits investigated, supergroup A infection show such negative effects only for a few of those traits. By testing for differential titer of bacteria by qRT-PCR, we also show a higher density of supergroup B and double infected strains across the different developmental stages of *N. vitripennis* males and females as compared to the supergroup A infected ones.

## Materials and Methods

### *N. vitripennis* strains used, their *Wolbachia* infections, and strain nomenclature

The *N. vitripennis* strains used NV PU-14 were obtained from Mohali, Punjab, India, in 2015. NV PU-14 was cured of *Wolbachia* by feeding the females with 1 mg/ml tetracycline in 10 mg/ml sucrose solution for at least two generations (Breeuwer & Werren, 1990). The curing was confirmed by PCR using supergroup-specific *ftsZ* primers (Baldo et al., 2006), and CI crosses between the infected and uninfected strains. NV PU-14 also served as the source strain for separating the two *Wolbachia* infections into single A and single B infected strains.

To separate the strains, we utilized relaxed selection on the females by repeatedly mating them with uninfected males which were obtained by antibiotic curing of the same NV PU-14 strain. Uninfected males do not have any sperm modification by *Wolbachia* which results in the removal of any selection pressure on the females to maintain their *Wolbachia* infections. Repeated mating with uninfected males was continued for ten generations till some of the progenies were found to be infected with either single A or single B supergroup infections. The single infection status of these strains was confirmed by using supergroup-specific *ftsZ* gene PCR primers (Baldo et al., 2006).

The preferred method of nomenclature of *Nasonia* strains and their *Wolbachia* infections includes information on supergroups as well as the host genotype. For example, [*w*NvitA *w*NvitB]V-PU14 indicates that the host species is *N. vitripennis*, with NV-PU14 as the host genotype, which has two *Wolbachia* infections, one each from supergroup A and supergroup B. However, since we used only *N. vitripennis* strains in this study, the nomenclature has been simplified by removing the species name. For example, the same strain will now be denoted as *w*A*w*B(PU), and when cured of these infections, as 0(PU). The single *Wolbachia*-infected strains used were designated as *w*A(PU) for the supergroup A-infected strain while *w*B(PU) for the supergroup B-infected strain. As the cured 0(PU) lines were in culture for three years, many of the infected lines were cured again to obtain ‘recently cured’ lines to minimize the effects of any host divergence that might have accumulated within them. These ‘recently cured’ lines were named 0(*w*A PU), 0(*w*B PU), and 0(*w*A*w*B PU). All these wasp strains were raised on *Sarcophaga dux* fly pupae with a generation time of 14-15 days at 25°C, 60% humidity, and a continuous daylight cycle.

### Sequential mating and sperm depletion of the males

To test the effect of *Wolbachia* on male reproductive traits like mating ability, individual males were assayed for the number of copulations it can perform, as well as sperm depletion. As *N. vitripennis* is haplodiploid, every successful mating will result in both female and male progenies while an unsuccessful one will result in the production of exclusively male progenies. The males used were obtained from virgin females hosted with one fly pupa for 24 hours and were not given any external sources of nutrition (usually a mixture of sucrose in water) before the experiment. Each male was then mated sequentially with virgin females from the same strain. At the first sign of a male not completing the entire mating behavior (Jachmann & Assem, 1996), it was given a rest for half an hour and was subjected to mating again until it stopped mating altogether. The mated females were hosted after a day with one fly pupa for 24 hours. The females were then removed and the offspring were allowed to emerge and then counted. The average number of copulations and the number of copulations before sperm depletion were compared using the Mann-Whitney U test with a significance level of 0.05.

### Host longevity, family size, and fecundity

To test whether the presence of *Wolbachia* has any influence on longevity, emerging wasps of both the sexes were kept individually in ria vials at 25°C, without any additional nutrition. Survival following emergence was measured by counting the number of dead individuals every 6 hours. Kaplan-Meier analysis followed by Log Rank Statistics was used to identify differences between strains with a significance level of 0.05.

To test for the effect of *Wolbachia* infections on the adult family size of virgin and mated females, each female was sorted at the pupal stage and separated into individual ria vials. To enumerate the brood size of mated females, some of these virgins were offered single males from the same strain and observed till mating was successful. All the females were then hosted individually with one fly pupa for 24 hours. These were kept at 25°C for the offspring to emerge which were later counted for family size by randomizing the ria vials in a double-blind assay. The differences were compared using the Mann-Whitney U test with a significance level of 0.05.

To investigate if *Wolbachia* affects the female fecundity, emerged females were hosted with one host for 24 hours. The host pupa was placed in a foam plug so that only the head portion of the pupa was exposed and available for the females to lay eggs. They were removed after 24 hours and the eggs laid were counted under a stereomicroscope (Leica M205 C). The differences in fecundity were compared using the Mann-Whitney U test with a significance level of 0.05.

### Estimation of relative density of Wolbachia infections across different developmental stages of *N. vitripennis*

To collect the different developmental stages, females were hosted for 4 hours, (instead of 24 hours in the previous experiments), with one host to narrow down the developmental stages of the broods. The larval and pupal stages (from Day 3 to Day 13 for males and from Day 8 to Day 14 for females) were collected every 24 hours. Larval stages for females were not done to avoid any DNA contamination from the males as the two sexes are virtually indistinguishable at these stages. Three replicates of ten larvae or pupae from the three strains, *w*A(PU), *w*B(PU), and *w*A*w*B(PU), were collected for each developmental stage. DNA extraction was done using the phenol-chloroform extraction method, where samples were crushed in 200 µl of 0.5 M Tris-EDTA buffer with 1% sodium dodecyl sulfate (SDS) and 2 µl of 22 mg/ml Proteinase K and incubated overnight at 37°C. DNA was purified with buffer saturated phenol and chloroform-isoamyl alcohol solution (24:1) and precipitated overnight with isopropanol at -20° C. The precipitated DNA pellet was dissolved in 60 µl nuclease-free water. The DNA concentration of the samples was measured using the Nanodrop 2000® spectrophotometer (Thermo Scientific). The concentrations of all the samples were normalized to 200 ng/µl across the different male and female developmental stages to be used for quantitative PCR. CFX96 C1000® Touch Real-time qRT-PCR machine (BioRad) was used to assay the relative density of *Wolbachia* across the strains. Amplification was done for the *Wolbachia* hcpA gene (Forward Primer: 5’-CTTCGCTCTGCTATATTTGCTGC-3’, Reverse Primer: 5’-CGAATAATCGCAACCGAACTG-3’). The primers were tested to amplify both the *Wolbachia* supergroup A and B strains. *Nasonia* S6K was used as the control gene (Bordenstein and Bordenstein, 2011). The total reaction volume contained 5 μl of iTaq Universal SYBR® Green supermix (BIO-RAD), 0.05 μl each of 10 μM of forward and reverse primers, and 200 ng of template DNA for a total volume of 10 μl for each reaction. Uninfected *N. vitripennis* DNA was used as negative control while DNase-free water was used as non-template control. Reaction conditions included an initial denaturation step of 95°C for 3 minutes followed by 39 cycles of 95°C for 10 seconds, annealing, and amplification at 52°C for 30 seconds for 40 cycles. All the reactions were performed in triplicates and included a melt curve to check for non-specific amplification. The relative density of *Wolbachia* was estimated by calculating the mean delta threshold cycle (ΔC_q_), using the formula:

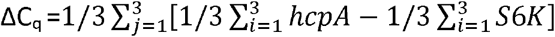

where i = number of technical replicates and j = number of biological replicates.

1/ ΔCq was calculated and plotted to show the *Wolbachia* density across different developmental stages of males and females. The relative *Wolbachia* density was compared across the three different strains using the Kruskal Wallis *H* test. Mann Whitney U test was used to compare two different groups with a significance level of 0.05.

## Results

### The presence of *Wolbachia* reduces the lifespan of both males and females

*Wolbachia* can compete with the host for available nutrition which can increase nutritional stress resulting in a shortened lifespan of many hosts (McMeniman et al., 2009; Caragata et al., 2014). Therefore, we first investigated the effect of *Wolbachia* infections on the survival of both male and female wasps. As Fig. 1(A), indicates, there is a significant difference in the longevity of the infected males across the three infection types. The double-infected strain, *w*A*w*B(PU), starts to die-off first and has a significantly shorter life span compared to the two single infected strains {Log Rank Test, χ^2^=16.8, p < 0.001 for *w*A(PU) and χ^2^=33.9, p < 0.001 for *w*B(PU)}. Males from the uninfected strain, 0(PU), lived the longest and showed significantly longer lifespan compared to all the other infected strains {Log Rank Test: χ^2^=76.3, p < 0.001 for *w*A*w*B(PU); χ^2^=33.0, p < 0.001 for *w*A(PU) and χ^2^=16.3, p < 0.001 for *w*B(PU)}. However, there was no significant difference in the lifespan of the two single infected strains *w*A(PU) and *w*B(PU) {Log Rank Test, χ^2^=3.84, p = 0.05}. Similarly, infected females (Fig. 1B) also showed a distinct reduction in life span when compared with the uninfected strain. But, unlike the males, the single A infected *w*A(PU) females, showed the shortest lifespan {Log Rank Test: χ^2^=11.2, p < 0.001 for *w*A*w*B(PU), χ^2^=56.9, p < 0.001 for *w*B(PU) and χ^2^=31.1, p < 0.001, for 0(PU)} followed by *w*A*w*B(PU) {Log Rank Test: χ^2^=20.4, p < 0.001 for *w*B(PU) and χ^2^=12.9, p < 0.001 for 0(PU)}. Curiously, 0(PU) and *w*B(PU) females showed similar lifespans {Log Rank Test: χ^2^=0.24, p = 0.62}. These results indicate a sex-specific variation as *w*A*w*B(PU) show the shortest lifespan among the males but *w*A(PU) showed the shortest among the females. Moreover, the effect of single infections on longevity also varied among the sexes as *w*A(PU) and *w*B(PU) males had similar lifespans, but in the case of females, it was *w*B(PU) and 0(PU) which had similar lifespans. But what is unambiguous from these results is that the uninfected strain lived the longest, irrespective of the sex of the hosts, indicating that the presence of *Wolbachia* is associated with the reduction in lifespan and is thus, costly for *N. vitripennis* to maintain.

**Figure 1.**
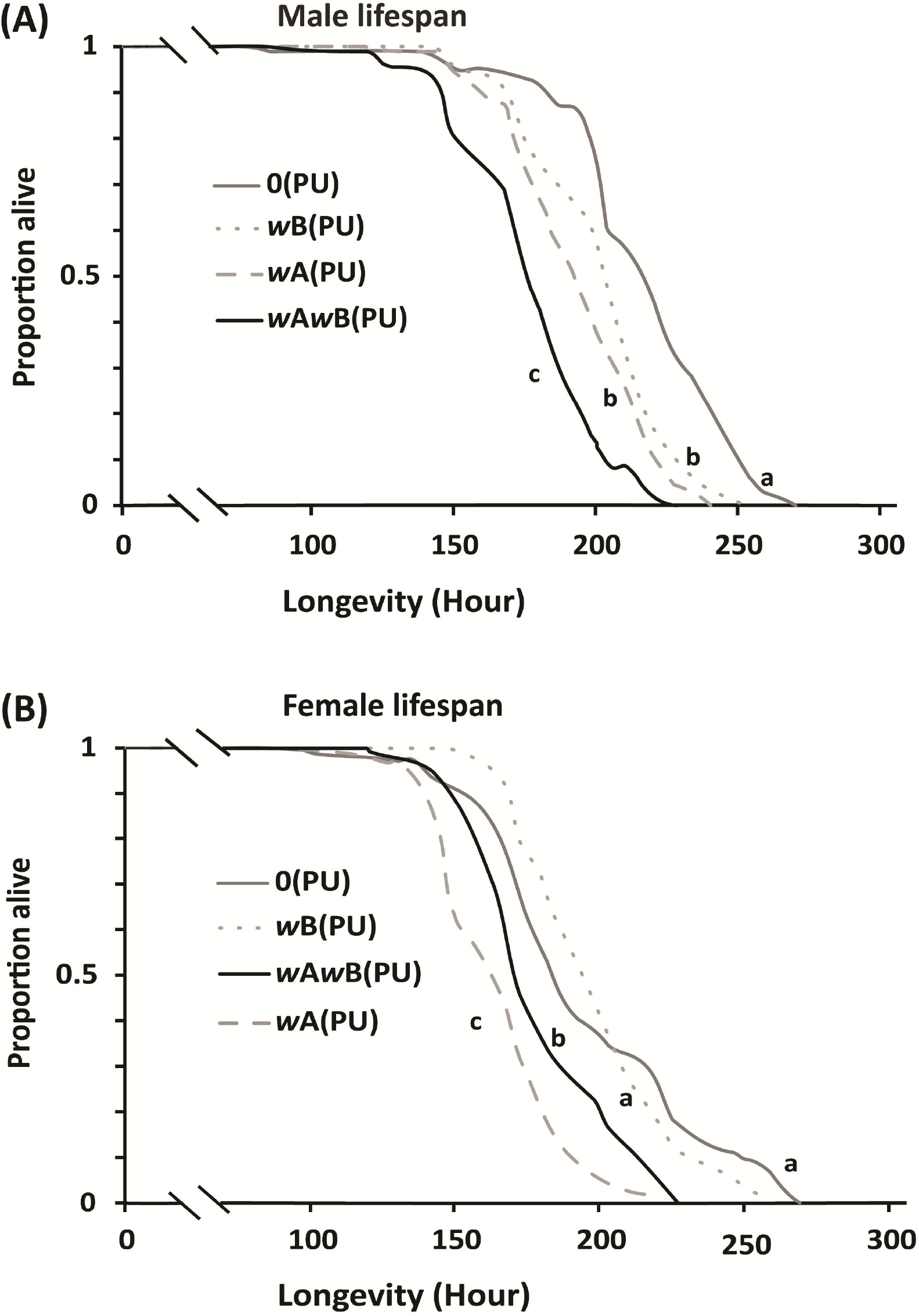
*Wolbachia* infected males and females show reduced lifespan. (A) : Lifespan of males. Sample sizes for the four infections 0(PU), *w*A(PU), *w*B(PU) and *w*A*w*B(PU) were n=98, n=95, n=94 and n=95 respectively. (B): Lifespan of females. Sample sizes for the four infections 0(PU), *w*A(PU), *w*B(PU) and *w*A*w*B(PU) were n=73, n=96, n=95 and n=95 respectively. Statistical significance was tested using Log Rank statistics with p < 0.05.

### The presence of *Wolbachia* reduces the number of copulations a male can perform

*Wolbachia* is known to be associated with a reduction in the number of mating a male can perform in *Ephestia kuehniella* (Sumida et al., 2017). To test whether similar effects are seen in *N. vitripennis*, we enumerated the number of copulations an individual male can perform across the infection types. As Fig. 2, indicates, there is indeed a reduction in the capacity of infected males to mate. When compared with the uninfected strain 0(PU), this reduction was most pronounced in *w*B(PU) (MWU, U=30, p < 0.01), followed by *w*A*w*B(PU) and *w*A(PU) which showed similar copulation capacities (MWU: U=11, p = 0.49). The uninfected 0(PU) strain produced males with the highest number of copulations {MWU: U= 32, p < 0.05 for *w*A(PU) and U= 27, p < 0.05 for *w*A*w*B(PU)}. Thus, the presence of *Wolbachia* substantially reduced the number of copulations that a male could perform. However, complex phenotypes like copulation can also be affected by the host genotype. Although all these four strains were derived from the same field-collected isofemale line, continuous culturing in the laboratory can fix specific alleles within them resulting in interstrain divergence. Moreover, it is also known that in *Nasonia* the effect of *Wolbachia*-induced phenotype is influenced by the hosts’ genetic background (Raychoudhury et al., 2012). Therefore, we cured all these infections again and tested whether the host genotype, rather than *Wolbachia*, is causing this reduction in the number of copulations by comparing these newly cured strains back with the previously used uninfected strain, 0(PU). As Fig. 2, indicates, males from most of these re-cured strains showed a marked and significant increase in the number of copulations performed. This number in the re-cured double infected strain, 0(*w*A*w*B PU), increased to similar levels as shown by 0(PU) (an increase of 29%, MWU: U=9.5, p = 0.2), while also showing a significant increase from its infected counterpart *w*A*w*B(PU) (from 73.5 ± 10.5 to 94.8 ± 15.39, MWU: U=3, p < 0.05). Similarly, the number of copulations for the re-cured single A supergroup infected strain, 0(*w*A PU), also increased to the levels of the uninfected strain 0(PU) (an increase of 7%, MWU: U=20, p = 0.76). However, this increase (from 77.5 ± 6.3 to 83.5 ± 12.9) with its infected counterpart was not significant (MWU: U=23, p = 0.48). The re-cured single B supergroup infection, 0(*w*B PU), was the only strain which did not revert to uninfected levels (MWU: U=22, p < 0.05) despite showing an increase (from 62.8 ± 6.6 to 78.2 ± 5.1; MWU: U=1, p < 0.05). However, what is evident is that the presence of *Wolbachia* is also associated with a reduction in the capability of a male to mate. Furthermore, by curing the infected strains again, we showed that this decrease is not due to the host genotype but is an effect of *Wolbachia* present in these strains.

**Figure 2.**
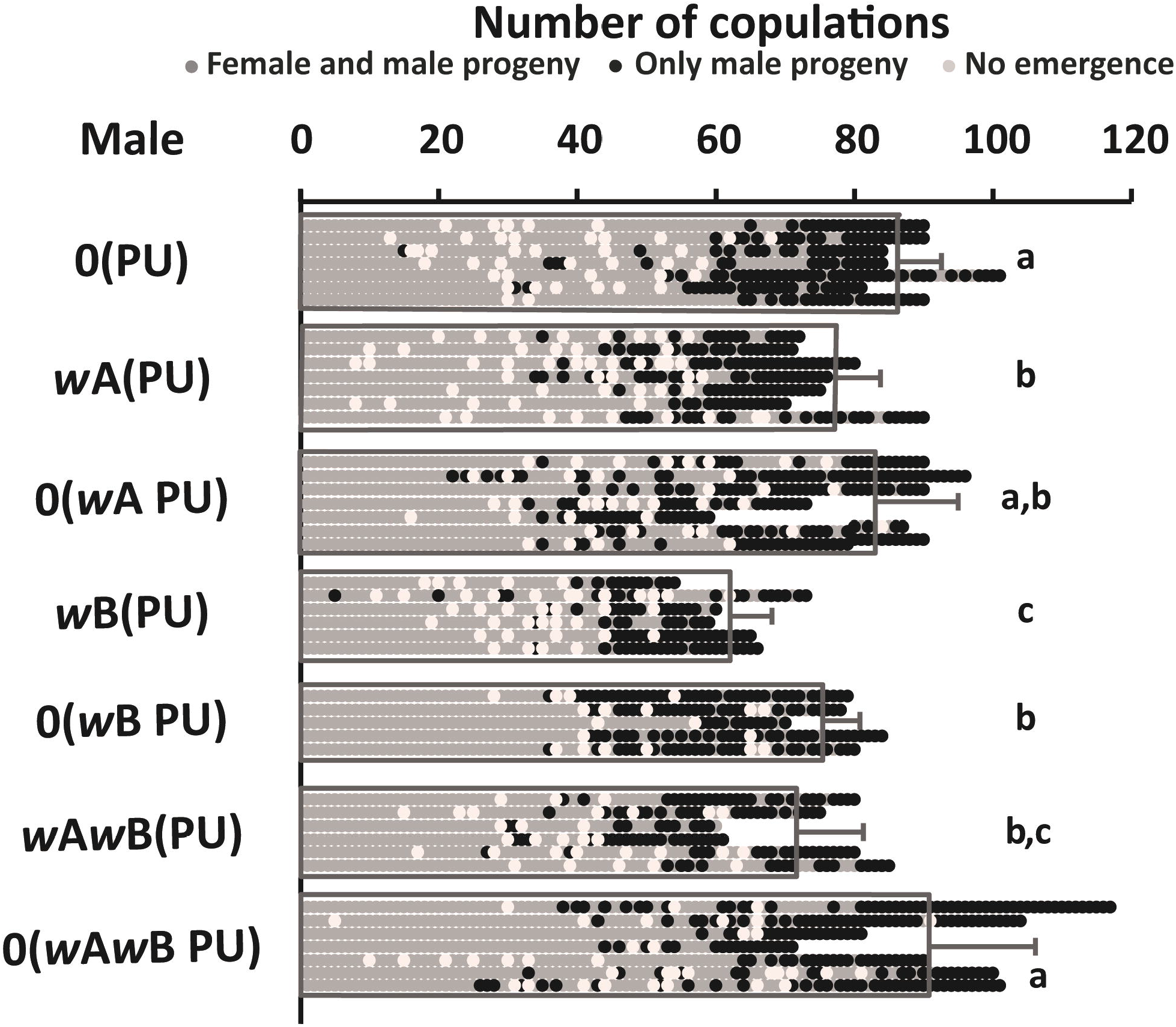
*Wolbachia* infected males show a reduction in the number of copulations. Males from different *Wolbachia* infections were mated sequentially until each of them stopped mating. 0(PU) could perform the highest number of copulations while *w*B(PU) the least. The results show that the presence of *Wolbachia* is associated with the reduction in the number of copulations a male can perform. The figure also shows whether the progenies of these sequential copulations produce any daughters or not, as a measure of sperm depletion. The details of sperm depletion are shown in figure 3. Sample sizes for the strains 0(PU), *w*A(PU), 0(*w*A PU), *w*B(PU), 0(*w*B PU), *w*A*w*B(PU) and 0(*w*A*w*B PU) were n=7, n=7, n=7, n=6, n=5, n=6 and n=7 respectively.

### *Wolbachia* infected males deplete their sperm reserves faster than the uninfected ones

*N. vitripennis* males are prospermatogenic (Boivin et al., 2005), where each male emerges with their full complement of mature sperm and have not been reported to produce any more during the rest of their lifespan (Chirault et al., 2016). Thus, if a single male is mated sequentially with as many females it can mate with, it should eventually run out of this full complement of sperm resulting in an all-male brood even after successful copulation. As Fig. 2, indicates, each male did run out of sperm at the tail end of the continuous mating and produced only male progenies (shown by black dots). We looked at the number of mating done by these males before sperm depletion to see if *Wolbachia* affects the sperm production in the males. As shown in Fig. 3, the average number of daughter progenies reduced with the number of mating (shown by the primary Y-axis on the left), indicating sperm depletion. Similar to copulation numbers, *w*B(PU) males were also the quickest to deplete their sperm reserve {MWU: U=27.5, p < 0.05 for *w*A(PU) and U=30, p < 0.01 compared for 0(PU)}. This was followed by *w*A*w*B(PU) and *w*A(PU) (MWU: U=13, p = 0.7). However, the uninfected males from 0(PU) were the slowest to deplete their sperm {MWU: U= 35, p < 0.01 for *w*A(PU) and U= 24, p < 0.05 for *w*A*w*B(PU)}. We again tested whether the host genotype, rather than *Wolbachia*, is causing this rate of sperm depletion, by comparing it with the recently cured strains. As shown in Fig. 3, the number of mating before sperm depletion increased for the recently cured 0(*w*A PU) up to the levels of 0(PU) (an increase of 5%, MWU: U=30, p = 0.06). However, this increase (from 48.14 ± 4.94 to 50.57 ± 9.41) was not significantly different from their infected counterpart *w*A(PU) (MWU: U=16, p = 0.8). Rates of depletion for 0(*w*A*w*B PU) also increased up to the levels of 0(PU) (an increase of 15.2%, MWU: U=21, p = 0.66). Again, the cured strain 0(*w*B PU), increased from *w*B(PU) (from 41 ± 1.67 to 47.6 ± 6.0,MWU: U=0, p < 0.05) but was still lower in number than 0(PU) (MWU: U=23, p < 0.05). These results indicate that the presence of *Wolbachia* has a significant negative impact on the number of sperm produced or utilized by the infected strains.

**Figure 3.**
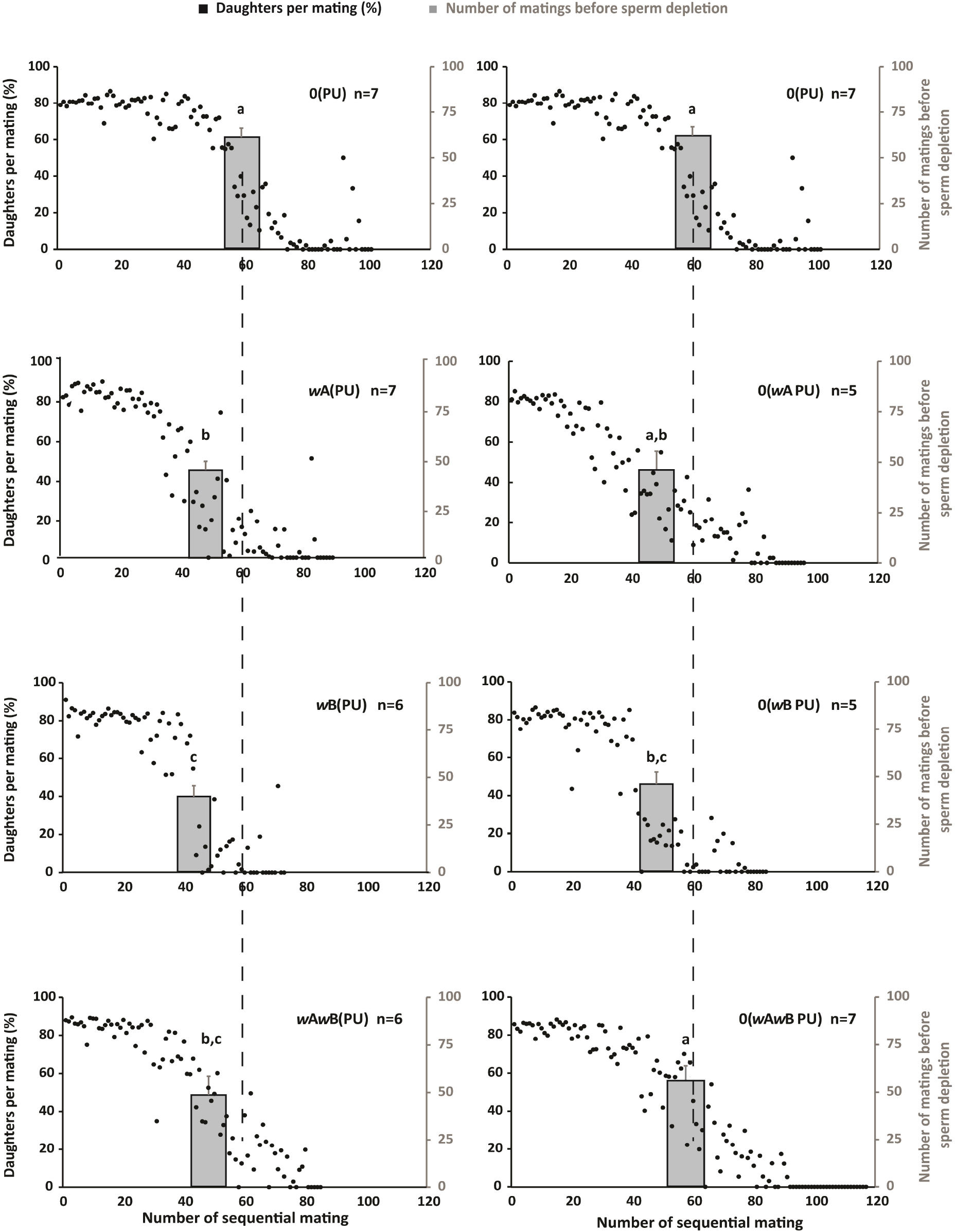
*Wolbachia* infected males deplete their sperm faster than the uninfected males. The Y-axis in black on the left of each figure represents the percentage of daughters produced for each mating. The black dots represent the average number of daughters produced for each sequential mating by the males of different *Wolbachia* infections (detailed in figure 2). The number of daughters produced is taken as a measure of the number of sperm transferred during each mating. The Y-axis, in grey, on the right, for each figure tallies the average number of copulations that yielded at least one daughter. Thus, it measures the number of mating before a male is depleted of its sperm. The left panel shows the males from *Wolbachia* infected strains whereas the right panel shows their respective cured versions. Data for 0(PU) is repeated at the top for comparison. The statistical significance was tested using the Mann-Whitney U test with p < 0.05.

### *Wolbachia* infected females produce fewer offspring

*Wolbachia* is known to have a negative impact on the progeny family size of its host (Hoffmann et al., 1990; Hohmann et al., 2001). To test whether a similar effect is seen in *N. vitripennis*, we enumerated the family sizes for both virgin and mated females for the four different *Wolbachia* infection strains and their recently cured counterparts.

As Fig. 4(A), indicates, there is a significant reduction in the average family sizes of all-male broods produced by the virgin females of the *Wolbachia* infected strains. When compared with the uninfected strain 0(PU), this reduction was most pronounced in *w*A*w*B(PU) (MWU: U=21151.5, p < 0.01) followed by *w*B(PU) (MWU: U=19880.5, p < 0.05). *w*B(PU) and *w*A*w*B(PU) showed similar family size (MWU: U=18582.5, p = 0.29). However, *w*A(PU) showed similar family size when compared with 0(PU) (MWU: U=17191, p = 0.39) and *w*B(PU) (MWU: U=17284, p = 0.26) but had larger all-male broods sizes than *w*A*w*B(PU) (MWU: U=18252, p < 0.05). We also compared the recently cured single and double infected strains with the infected parental strains. The recently cured strains 0(*w*B PU) and 0(*w*A*w*B PU) showed marginal increase in the family size which was comparable to the uninfected strain 0(PU) {an increase of 1.5%, MWU: U=11554, p = 0.29 for 0(*w*B PU); an increase of 2%, MWU: U=10798, p = 0.21 for 0(*w*A*w*B PU)}. However this marginal increase {from 29.84 ± 12.94 to 31.44 ± 10.30 for 0(*w*B PU) and from 28.99 ± 11.60 to 30.97 ± 11.48 for 0(*w*A*w*B PU)} was not significantly different from their infected counterparts *w*B(PU) (MWU: U=9963.5, p = 0.34) and *w*A*w*B(PU) (MWU: U=8650.5, p = 0.09). The recently cured strain 0(*w*A PU) did not show any increase in the family size and was comparable to *w*A(PU) (MWU: U=7085.3, p = 0.63) and 0(PU) (MWU: U=8161, p = 0.22).

**Figure 4.**
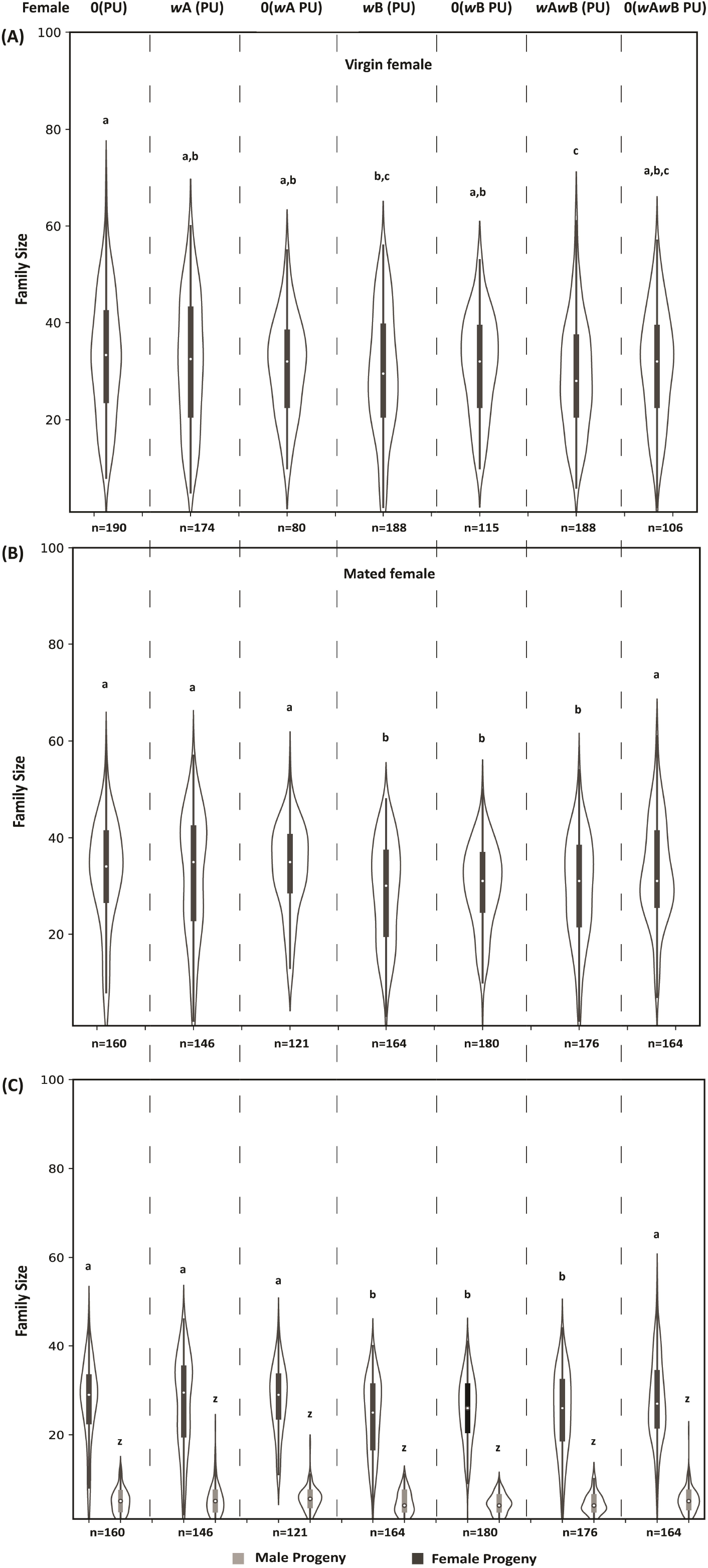
*Wolbachia* infected females produce fewer offspring. Family size produced by females when hosted as virgins (A) and mated (B). The difference in the family size of mated females is due to the difference in the number of daughters (C) as there is no significant difference in the number of males produced. The statistical significance is tested using the Mann-Whitney U test with p < 0.05.

Similar to the virgin females, the reduction was also observed for average family sizes of mated females as shown in Fig. 4(B). When compared with the uninfected strain 0(PU), this reduction is most pronounced in *w*B(PU) (MWU: 15582, p < 0.01) and *w*A*w*B(PU) (MWU: U=16303, p < 0.01). However, *w*B(PU) and *w*A*w*B(PU) show similar levels of family sizes (MWU: 13732.5, p = 0.55). Interestingly, *w*A(PU) strain shows similar levels of family size as 0(PU) (MWU: U=11396.5, p = 0.86) but had larger family sizes than *w*B(PU) (MWU: U=14080, p < 0.01) and *w*A*w*B(PU) (MWU: U=14682, p < 0.05). Upon curing, the average family sizes of the recently cured 0(*w*A*w*B PU) reverted back to the levels of the uninfected strain 0(PU) (an increase of 11.8%, MWU: U= 13295, p = 0.61) showing a significant increase from the infected counterpart *w*A*w*B(PU) (from 29.49 ± 10.67, MWU: U=12023, p < 0.05). The recently cured strain 0(*w*A PU) did not show any significant increase from the infected counterpart *w*A(PU) (MWU: U=8385.5 p = 0.69) and was comparable to 0(PU) (MWU: U=9022.5, p = 0.5). However, the recently cured strain 0(*w*B PU) did not increase to the levels of the uninfected strain 0(PU) (an increase of 4.3%, MWU: U=16782, p < 0.05). The marginal increase in the family size (from 28.72 ± 10.46 to 29.97 ± 8.59) was not significantly different from the parent strain *w*B(PU) (MWU: U=13854, p = 0.47).

To understand whether this difference in the family size of the mated females is due to the production of fewer daughters or sons or both, we compared their numbers separately for the four strains (Fig. 4C). No difference was observed in the number of sons produced by the mated females. However, significant differences were observed in the number of daughters produced. When compared to the uninfected strain 0(PU), *w*B(PU) and *w*A*w*B(PU) showed the least number of daughters produced {MWU: 15964, p < 0.01 for *w*B(PU) and U=16283, p < 0.01 for *w*A*w*B(PU)} whereas *w*B(PU) and *w*A*w*B(PU) produced nearly equal number of daughters (MWU: 13392, p = 0.33). Again, *w*A(PU) strain produced equal number of daughters compared to 0(PU) (MWU: U=11543, p = 0.98) but higher in number than *w*B(PU) and *w*A*w*B(PU) {MWU: U=14201, p < 0.01 for *w*B(PU) and MWU: U=14372, p < 0.05 for *w*A*w*B(PU)}. Upon curing, the recently cured 0(*w*A*w*B PU) reverted to the levels of the uninfected strain 0(PU) (MWU: U= 13545, p = 0.42) showing a significant increase in the number of daughters from the infected counterpart *w*A*w*B(PU) (MWU: U=12331, p < 0.039). The recently cured strain 0(*w*A PU) did not show any increase in the number of daughters produced from their infected counterpart *w*A(PU) (MWU: U=8468 p = 0.79) and was also comparable to 0(PU) (MWU: U=9330, p = 0.84). However, recently cured strain 0(*w*B PU) did not increase to the levels of the uninfected strain 0(PU) (MWU: U=16749.5, p < 0.01).

### *Wolbachia* negatively impacts the fecundity of infected females

To check whether the differences in the family sizes between the different infection strains are due to the number of eggs being laid by the females, we looked at the fecundity of both virgin and mated females across the strains. Among the virgin females (Fig. 5A), *w*A*w*B(PU) had the least fecundity (MWU: U=5042, p < 0.01) when compared to the uninfected strain 0(PU). However, there was no significant difference between 0(PU) and *w*A(PU) (MWU: U=2632.5, p = 0.34) and also between 0(PU) and *w*B(PU) (MWU: U=2052.5, p = 0.71).

**Figure 5.**
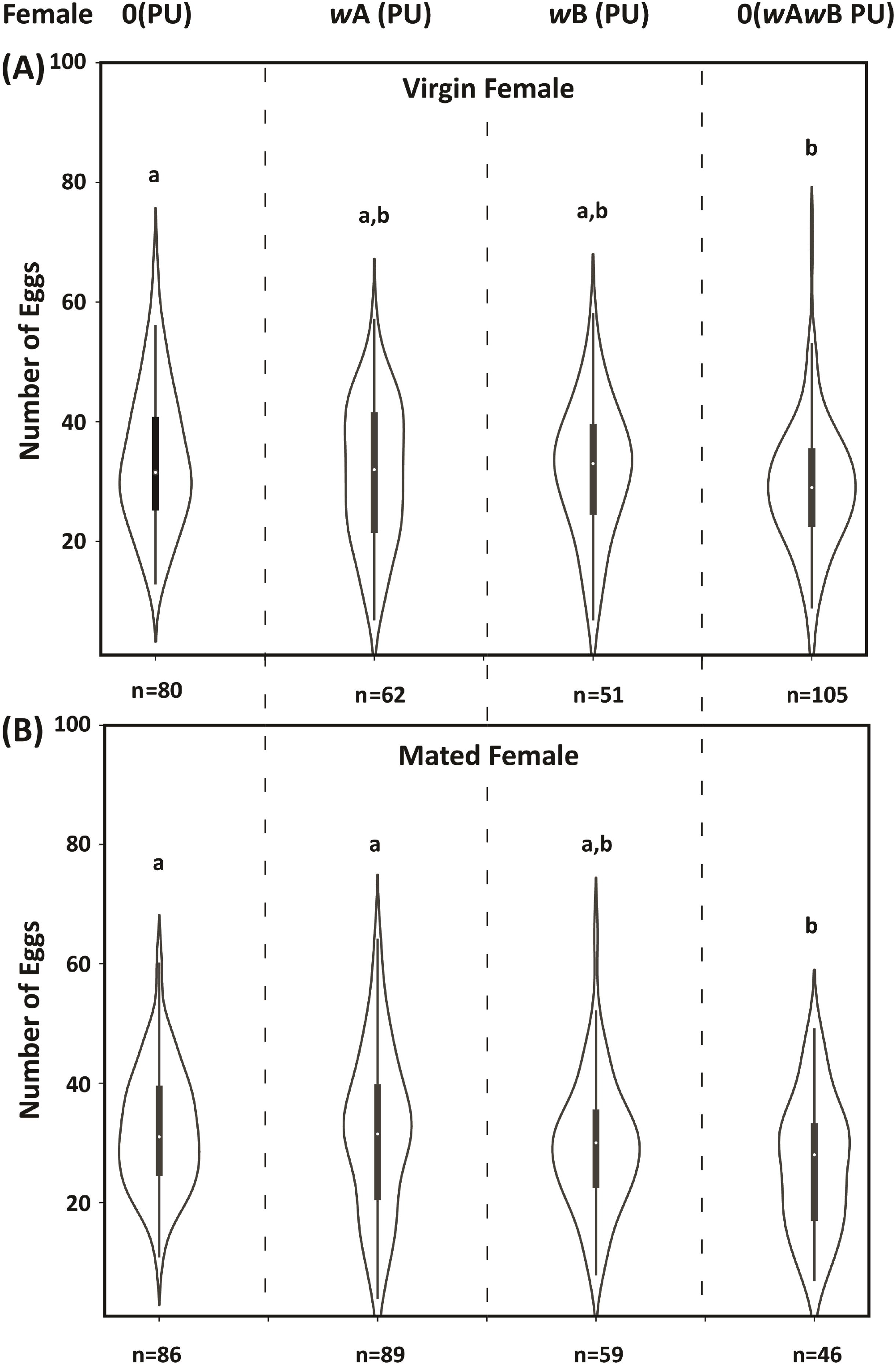
*Wolbachia* infection reduces female fecundity. The measure of fecundity (number of eggs laid) by females of different infections, with virgin females (A) and mated females (B). For virgin females, 0(PU) shows higher fecundity than *w*A*w*B(PU), while *w*A(PU) and *w*B(PU) strains have intermediate fecundity which is comparable to both 0(PU) and *w*A*w*B(PU). However, for mated females, 0(PU) and *w*A(PU) show higher fecundity than *w*B(PU) and *w*A*w*B(PU). The statistical significance is tested with the Mann-Whitney U test, p<0.05.

Among the mated females (Fig. 5B), *w*A*w*B(PU) again had the least fecundity (MWU: U=2461.5, p < 0.01) when compared to the uninfected strain 0(PU). This was followed by *w*B(PU) having similar fecundity as that of *w*A*w*B(PU) (MWU: U=1479.5, p = 0.24) and *w*A(PU) (MWU: U=2822.5, p = 0.22). However, *w*A(PU) showed higher fecundity than *w*A*w*B(PU) (MWU: U=2388, p < 0.05) and was similar to the uninfected strain 0(PU) (MWU: U=3799.5, p = 0.75).

The double infected strain showed the least fecundity, both for the virgin and mated females, suggesting an effect of *Wolbachia* on egg production in females. The assay also established that the difference in family sizes is due to the differences in the fecundity of the females.

### Relative *Wolbachia* density in single and multiple *Wolbachia* infection strains

*Wolbachia* density has a major role to play in expressing the effects of the infection on host biology (Hoffmann et al., 1996; Min & Benzer, 1997). An increase in cellular *Wolbachia* density is often associated with a greater expression of their effects (Breeuwer & Werren, 1993). Thus, we estimated *Wolbachia* titers across different developmental stages of *N. vitripennis* males and females. In the case of males (Fig. 6A) *w*A(PU) had the lowest *Wolbachia* density across different larval and pupal developmental stages when compared with *w*B(PU) (MWU: U=11, p < 0.01) and *w*A*w*B(PU) (MWU: U=12, p < 0.01) This difference was more prominent at the adult stage. However, no such differences were found between *w*B(PU) and *w*A*w*B(PU) (MWU: U=51, p = 0.56).

**Figure 6.**
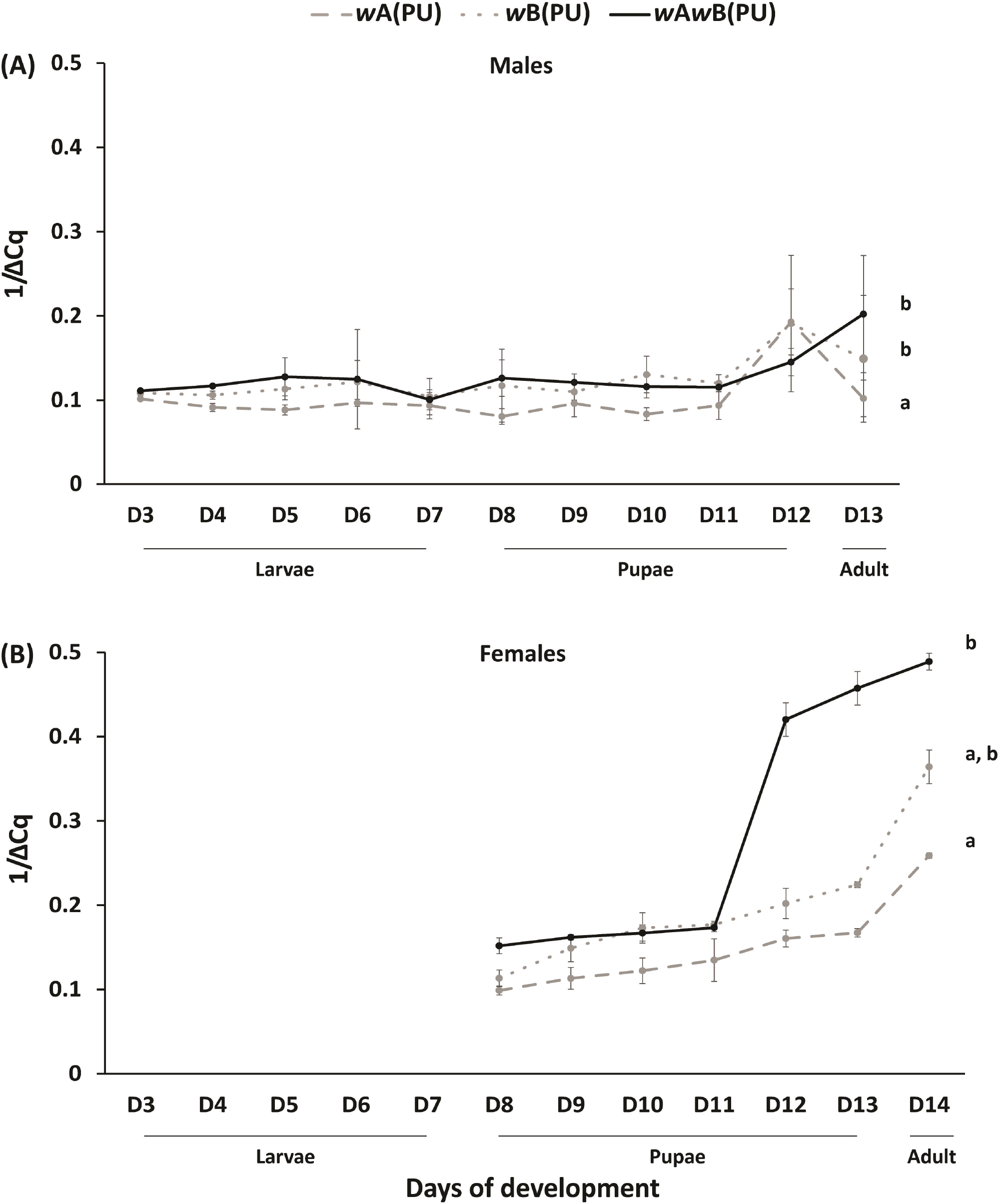
Quantitative estimation of *Wolbachia* across different developmental stages of *N. vitripennis* males (A) and females (B). The statistical significance between groups is tested using the Mann-Whitney U test, p < 0.05.

In the case of females (Fig. 6B), *w*A(PU) showed lower levels of *Wolbachia* when compared to *w*A*w*B(PU) (MWU: U=8, p < 0.05) again at the late pupal and adult stage. However, no difference was observed between *w*A(PU) and *w*B(PU) (MWU: U=12, p = 0.12) and also *w*B(PU) and *w*A*w*B(PU) (MWU: U=19, p = 0.5).

## Discussion

CI-inducing *Wolbachia* are known to have negative effects on various physiological traits in the vast majority of its host population (summarized in Table 1). The present study also suggests such effects, or a “cost”, associated with the maintenance of *Wolbachia* infection in *N. vitripennis*. This is in contrast to the previous reports suggesting positive fitness effects (Stolk C & Stouthamer R, 1996) and no fitness effects (Bordenstein & Werren, 2000) of *Wolbachia* on *N. vitripennis*. However, several unique conclusions emerge from the study.

Our results (summarized in Table 2) demonstrate a sex-independent cost of the presence of single and multiple *Wolbachia* infections. Many phenotypes show reduction across the sexes such as longevity (Fig. 1) where infected males and females showed reduced lifespan. When compared with the uninfected strain 0(PU), the *Wolbachia* infected strains *w*B(PU) and *w*A*w*B(PU) have reduced life spans. However, sex-specific variations were also observed among the infected strains where *w*A(PU), in females, had the shortest life span, while in the case of males, had a greater life span than *w*A*w*B(PU).

**Table 2.** Effect of *Wolbachia* infections on *N. vitripennis* (Summary)

| Phenotype | Host Sex | Effect of *Wolbachia* | "Cost" compared to 0(PU) |
| --- | --- | --- | --- |
| Lifespan | Male | $0(\text{PU}) > wB(\text{PU}) = wA(\text{PU}) > wAwB(\text{PU})$ | $wA(\text{PU}) = 11.1\%$, $wB(\text{PU}) = 6.5\%$,<br>$wAwB(\text{PU}) = 15.5\%$ |
| | Female | $0(\text{PU}) = wB(\text{PU}) > wAwB(\text{PU}) > wA(\text{PU})$ | $wA(\text{PU}) = 17.7\%$, $wAwB(\text{PU}) = 15.5\%$ |
| Number of copulations | Male | $0(\text{PU}) > wA(\text{PU})$ $wB(\text{PU})$ and $wAwB(\text{PU})$<br>$wA(\text{PU}) = wAwB(\text{PU})$, $wA(\text{PU}) > wB(\text{PU})$,<br>$wAwB(\text{PU}) = wB(\text{PU})$ | $wA(\text{PU}) = 12.4\%$, $wB(\text{PU}) = 28.8\%$,<br>$wAwB(\text{PU}) = 16.7\%$ |
| Sperm depletion | Male | $0(\text{PU}) > wA(\text{PU})$ $wB(\text{PU})$ and $wAwB(\text{PU})$<br>$wA(\text{PU}) = wAwB(\text{PU})$, $wA(\text{PU}) > wB(\text{PU})$,<br>$wAwB(\text{PU}) = wB(\text{PU})$ | $wA(\text{PU}) = 19.4\%$, $wB(\text{PU}) = 31.3\%$,<br>$wAwB(\text{PU}) = 15.7\%$ |
| Progeny family size | Female |  |  |
| | a. Virgin | $0(\text{PU}) > wB(\text{PU})$ and $wAwB(\text{PU})$, $0(\text{PU}) = wA(\text{PU})$,<br>$wA(\text{PU}) > wAwB(\text{PU})$, $wA(\text{PU}) = wB(\text{PU})$,<br>$wB(\text{PU}) = wAwB(\text{PU})$ | $wB(\text{PU}) = 10\%$, $wAwB(\text{PU}) = 12.4\%$ |
| | b. Mated | $0(\text{PU}) = wA(\text{PU}) > wAwB(\text{PU}) = wB(\text{PU})$ | $wB(\text{PU}) = 11.5\%$, $wAwB(\text{PU}) = 9.1\%$ |
| Fecundity | Female |  |  |
| | a. Virgin | $0(\text{PU}) = wA(\text{PU}) = wB(\text{PU})$, $0(\text{PU}) > wAwB(\text{PU})$ | $wAwB(\text{PU}) = 14.1\%$ |
| | b. Mated | $0(\text{PU}) = wA(\text{PU}) > wB(\text{PU}) = wAwB(\text{PU})$ | $wB(\text{PU}) = 9.1\%$, $wAwB(\text{PU}) = 18.2\%$ |
| *Wolbachia* density | Male | $wAwB(\text{PU}) > wB(\text{PU}) = wA(\text{PU})$ | |
| | Female | $wAwB(\text{PU}) > wA(\text{PU}) = wB(\text{PU})$ | |

*Wolbachia* affected the reproductive capabilities of the infected males and reduced their copulation capability (Fig. 2) and also led to quicker sperm depletion (Fig. 3). Such negative effects on reproductive traits were also observed in females where the infected females produced fewer progenies (Fig. 4) compared to 0(PU). These differences were elicited at the level of female fecundity where the number of eggs being laid by the infected females was fewer (Fig. 5) indicating that the negative effects of *Wolbachia* manifested themselves even before the egg-laying stage. However, the egg to larval to pupal stage mortality could also have an effect on the brood sizes but these were not assayed.

In most cases, these negative effects disappeared with the removal of *Wolbachia*, indicating the role of *Wolbachia* in producing these negative effects and not the host genotype. In phenotypes like family sizes, the recently cured strains showed a significant increase suggesting that the negative effects are due to the presence of *Wolbachia*. However, 0(*w*B PU) did not revert to the levels of 0(PU) in the number of copulations performed and sperm depletion assays. A possible reason can be some residual effects of the parent genotype in 0(*w*B PU) males.

Our results also demonstrate supergroup-specific negative effects on the host. Supergroup B *Wolbachia* is costlier to maintain in both the sexes than supergroup A. While *w*B(PU) shows strong effects on all the traits studied across the sexes, *w*A(PU) had significant negative effects only on the reproductive traits of the males and the longevity of females. *w*A(PU) females, as an exception, had their family sizes comparable to 0(PU). These observations are unique as no comprehensive data is available on the supergroup-specific cost of *Wolbachia* infections.

The higher cost of maintenance of supergroup B *Wolbachia* can be an attribute of the CI phenotype induced by supergroup B *Wolbachia*. Complete CI (i.e, nearly 100% CI) are rare events reported mainly for supergroup B *Wolbachia* in *Culex pipiens, Aedes aegypti* (Sinkins et al., 2005; Xi et al., 2005), and in *N. vitripennis* as well (supplementary Fig. 1 and Bordenstein et al., 2006a). This essentially means that nearly the entire sperm complement of each male has the *Wolbachia*-induced CI modification and correspondingly, nearly all the eggs from the females have the rescue effect (Werren et al., 2008). Introducing 100% modification and rescue would necessitate relatively high *Wolbachia* titers to be maintained in both sexes which in turn can cause an elevated nutritional burden, eventually resulting in negative effects on the physiological traits of the host.

Previous reports have suggested a direct correlation between *Wolbachia* density and the level of CI (Breeuwer & Werren, 1993; Noda et al., 2001; Ikeda et al., 2003; Dutton & Sinkins, 2004; Ruang-Areerate & Kittayapong, 2006). Our results also suggest that the cost of *Wolbachia* maintenance is correlated with the density of *Wolbachia* strains present in the host. *Wolbachia* density in both the sexes differs in the late pupal stages (Day 11, Day 12) and adult stages. Thus, in the case of females, *w*A*w*B(PU), which shows a high bacterial load, has reduced fecundity and longevity. Similarly, in the case of males, the *w*A*w*B(PU) strain shows a reduced number of copulations and the number of sperm produced/transferred. *w*B(PU), therefore causes substantial detrimental effects on both the sexes of the host which is similar to the levels found in *w*A*w*B(PU). This again can be explained by the high *Wolbachia* load in both sexes. In contrast, *w*A(PU) showed minimal effect on the female hosts, with the longevity of females being an exception. A possible explanation for this can be the relatively low density of *w*A(PU) across the different developmental stages (Fig. 6) as compared to the other infections. Moreover, supergroup A infected strains are known to have relatively higher levels of phage density (Bordenstein et al., 2006). According to this phage density model, higher phage density has an impact on the level of CI caused by supergroup A strains, where it leads to a significant reduction in *Wolbachia* titer and hence has a reduction effect on CI phenotype. Our results also conform with the previous reports of the positive correlation between *Wolbachia* abundance and the level of CI induced not only in *N. vitripennis* (Bordenstein et al., 2006b) but in other insect taxa as well (Ijichi et al., 2002; Kondo et al., 2002). *w*B(PU) *Wolbachia* showed complete CI and had higher *Wolbachia* titers in both males and females, which was also comparable to that of the *w*A*w*B(PU), whereas *w*A(PU) had the lowest *Wolbachia* titers among the three strains and showed incomplete CI (supplementary Fig. 1). Thus, higher levels of *w*B(PU) than *w*A(PU) also explain the more severe effects of *w*B(PU) than *w*A(PU).

The low levels of *w*A(PU) and the associated negative effects on host physiology raise questions on the maintenance of *w*A(PU) infection in *N. vitripennis*. Since supergroup B appears to be a “stronger” *Wolbachia* than supergroup A, any competition for nutritional resources and niche habitation between them should drive out supergroup A *Wolbachia* from the hosts. But *w*A(PU) is observed to have less severe effects on the females with longevity being the only factor to have any negative effects. Supergroup A *Wolbachia* in *N. vitripennis* is closely related to other A supergroup *Wolbachia* like *w*Mel in *D. melanogaster* and *w*Ha, *w*Au, and *w*Ri in *D. simulans* (Díaz-Nieto et al., 2021). The presence of supergroup A *Wolbachia* in *N. vitripennis* could be due to such defenses against viral infections (Teixeira et al., 2008; Bhattacharya et al., 2017; Pimentel et al., 2021) but remain untested.

The varied effects of *Wolbachia* can be dependent on host strain as well as on a particular *Wolbachia* supergroup. Our data suggest that supergroup B *Wolbachia* behaves more like a parasite in both the sexes while supergroup A *Wolbachia* had detrimental effects mostly in the case of males. This is contrary to what is expected as the two *Wolbachia* supergroups being maternally transmitted, should affect the males more in comparison to the females as males are a dead-end for *Wolbachia* transmission.

Our experiments indicate an additive or synergistic effect of the presence of the two different *Wolbachia* supergroups in the double infected strain *w*A*w*B(PU). Evidence of such effects can be seen in traits like male longevity (additive effect) where the deficit in longevity for *w*A*w*B(PU) is equal to the total deficits caused by *w*A(PU) and *w*B(PU). Similarly, for traits like female longevity, virgin female family size, and female fecundity, the negative effects on strain *w*A*w*B(PU) appear to be a combined effect of *w*A(PU) and *w*B(PU) (synergistic effect). Since the two supergroup infections are bidirectionally incompatible with each other, it is plausible that they are also competing for the host nutrition, which can further enhance the negative impacts of these infections.

The negative fitness effects of CI-inducing *Wolbachia*, and nutritional competition raises important questions on the maintenance of these endosymbionts over long evolutionary time scales. Theoretical studies predict that for the long-term persistence of these maternally inherited reproductive parasites, they can eventually evolve towards mutualism (Prout, 1994; Turelli, 1994). Various theories on endosymbiosis also support the vertical transmission of endosymbionts, as the spread of the symbiont depends on the reproductive success of the host itself (Anderson & May 1982; Ewald, 1987; Bull et al., 1991; Lipsitch et al., 1995). Hence, mutualism can be a preferred way for endosymbionts to ensure their persistence. Moreover, if there are indeed some adverse effects of maintaining *Wolbachia*, then hosts would be under strong selection pressure to develop immunity against them. Evidence suggests that there are examples of such emergence of host genetic factors upon *Wolbachia* infection in hosts *Drosophila* and mosquitoes (Zug & Hammerstein, 2015). Host suppressor alleles have been identified conferring resistance against feminizing (Rigaud et al., 1999) and male-killing *Wolbachia* (Hornett et al., 2006). However, no such host genetic factors have been found for CI-inducing *Wolbachia*, especially in *N. vitripennis*. A possible explanation for the maintenance of these multiple infections comes from the high efficiency of transmission of these infections in *N. vitripennis*, where these infections also show nearly 100% transmission as well as CI (Breeuwer and Werren, 1990). For many *Wolbachia*-arthropod associations, the pathogenicity and transmission efficiency is mediated by bacterial titers (Correa & Ballard, 2016). Our results also show higher *Wolbachia* density for double infected strain, which can facilitate its transmission and maintenance in the host. Another possibility can be that these *Wolbachia* infections in *N. vitripennis* are relatively recent, the evidence of which comes from the rapid spread of *Wolbachia* in populations of *N. vitripennis* across North America and Europe (Raychoudhury et al., 2010b). These recent infections, although bearing a cost on the host at present, might eventually lead to the host immunity against the infection.

## Supporting information

Figure S1. CI crosses for single Wolbachia infection strains wA(PU) and wB(PU)

## Acknowledgments

We thank the Indian Institute of Science Education and Research (IISER) Mohali for the funding and graduate fellowship for AT. Partial funding was obtained from Grant No. BT/PR14278/BRB/10/1417/2015, Department of Biotechnology, Government of India, awarded to Nagaraj Guru Prasad (Associate Professor, Biological Sciences, IISER Mohali) and RR.

## Author Contribution

AT conceived the idea and planned the study. AT performed the experiments. AT collected the data. RB performed the pilot experiments for female fecundity. AT analyzed the data. AT, RS, and RR were involved in data representation and statistical analysis. AT and RR wrote the manuscript.

## Data Archiving

The raw data for all the experiments have been archived at Dryad with DOI: 10.5061/dryad.w0vt4b8s9

URL: https://datadryad.org/stash/share/gKvBkEcYjmrNULX4F53Ktu5Pt-P1pcXBnbA5Ib_HEPo

