## Supplementary material for "Bacterial supergroup specific “Cost” of *Wolbachia* infections in *Nasonia vitripennis*": Figure S1. CI crosses for single Wolbachia infection strains wA(PU) and wB(PU)

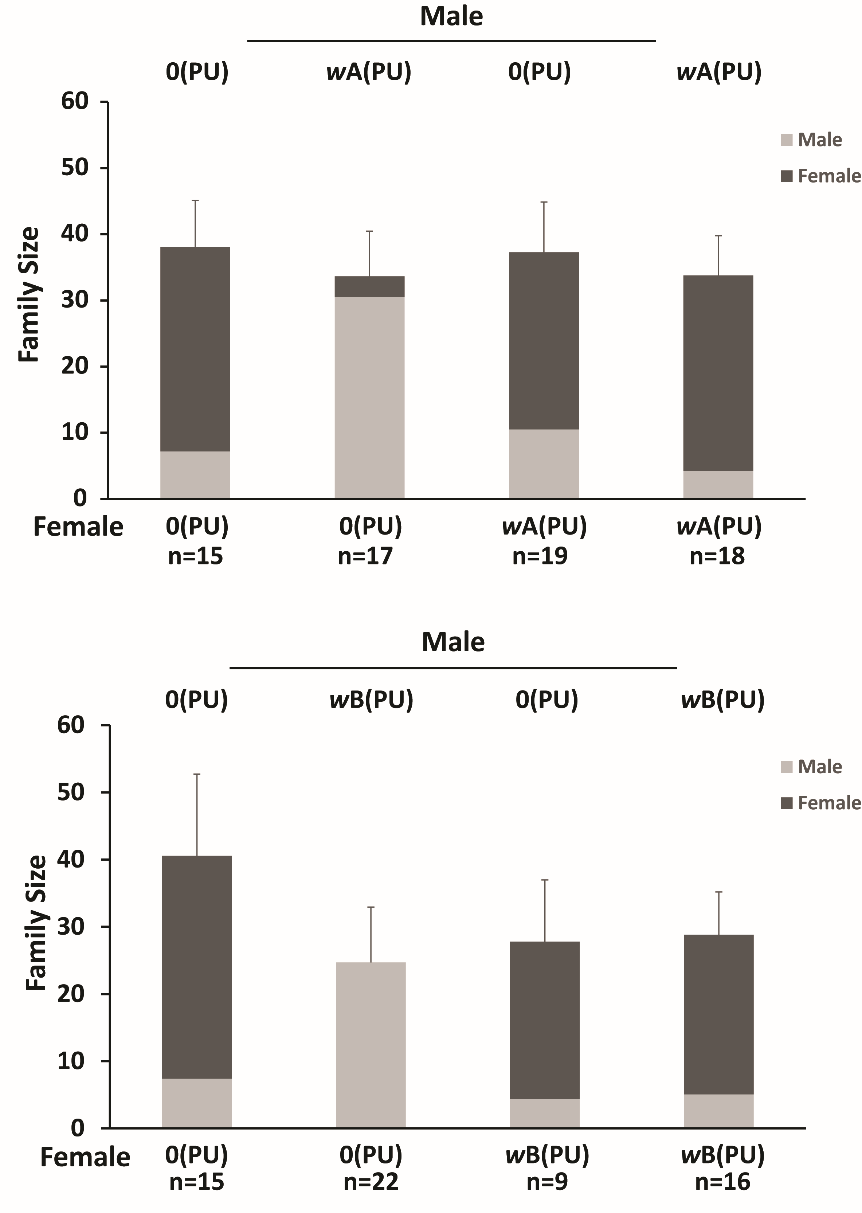

**Figure S1. CI crosses for single *Wolbachia* infection strains *w*A(PU) and *w*B(PU)**

*w*A(PU) males show incomplete CI with 0(PU) females as the cross produces female progenies as well (top panel). *w*B(PU) males show complete CI with 0(PU) females. The cross leads to the production of an all-male brood (bottom panel).
